# Diet-Induced Obesity is Associated with Sex-Dependent Cerebral Microvascular Changes and Cognitive Decline in Mice

**DOI:** 10.64898/2026.11.19.767808

**Authors:** L. Sheppard, S. Sawiak, C. Riches, S. Lundh, A. Secher, E. Nishimura, L.-S. Martis, M. Evans

## Abstract

Metabolic diseases, including obesity and diabetes, are major risk factors for cognitive impairment and dementia, yet the underlying causal mechanisms remain unclear. Here we investigate whether this link is mediated by dysfunctional changes in cerebral microvessels. Using a multimodal longitudinal approach combining 3D light-sheet fluorescence microscopy, contrast-enhanced MRI and cognitive testing, we characterise structural, functional and behavioural changes to the brain microvasculature in a mixed-sex mouse model of diet-induced obesity (DIO).

Our results reveal a striking sex-specific vulnerability to obesity-driven cerebrovascular dysfunction. Brains from male DIO mice exhibited reduced capillary diameter and volume in the frontal cortex, alongside decreased blood volume and accelerated cognitive decline over time in an object recognition task. Conversely, female DIO mice displayed compensatory capillary vasodilation and were protected against neurovascular and cognitive impairment. These findings link cerebral microvascular changes to obesity-associated cognitive decline, and identify sex-dependent differences in the cerebrovascular response to obesity.

## Main

Obesity and associated metabolic diseases, including diabetes, represent a growing global public health burden. In addition to their cardiovascular comorbidities, obesity and diabetes are strongly associated with cognitive impairment and dementia, including Alzheimer’s disease (AD)^1,2^. Despite a well-established association, the precise causal mechanisms linking metabolic disease to cognitive decline and dementia are not well understood. Cerebrovascular alterations may contribute to this association. Understanding how metabolic disease impairs cerebrovascular health is therefore critical for mitigating the growing burden of dementia in an increasingly metabolically compromised population.

Given that the brain requires a constant and reliable cerebral blood flow (CBF) for normal daily function, utilising a network of almost 600 km of capillaries, and receiving 20% of the body’s resting cardiac output ^3,4^, we hypothesize that obesity results in changes in the brain vasculature which adversely impact on brain function. A predominant mediator of vascular dysfunction is microvascular rarefaction, characterised by a decrease in the density of small blood vessels ^5^. Peripheral microvascular rarefaction occurs in metabolic disease in multiple tissues in both preclinical and clinical studies^6–10^. Moreover, cerebral microvascular rarefaction has separately been linked to neurodegeneration and vascular brain diseases including cerebral small vessel disease (cSVD), vascular dementia (VaD) and AD ^3,11,12^. Single-nucleus RNA sequencing analyses have further highlighted the enrichment of AD risk genes in human vascular cells from the frontal cortex and hippocampus ^13^. In addition to structural (i.e. anatomical) changes in blood vessels, microvessel changes may be functional, presenting as impaired or altered vessel perfusion ^14,15^. However, how obesity-associated changes in the brain microvascular structure relate to vascular function and cognitive decline, and whether these relationships differ between sexes, remains unclear.

To answer these questions, we combined whole-brain 3D light-sheet fluorescent microscopy (3D LSFM) and AI filament tracing with longitudinal ultra-high field MRI and serial cognitive testing to define the cerebrovascular consequences of diet-induced obesity (DIO) in mice, at a resolution previously unseen. Importantly, both sexes were included in the model, since there are sex differences in both metabolic disease and VaD disease risk, vascular risk profiles and biomarkers of cognitive impairment^16–18^.

Our findings reveal a male-specific vulnerability, with male but not female DIO mice exhibiting structural microvascular rarefaction in the frontal cortex. We found that alongside evidence of structural vascular changes, male DIO mice displayed signs of a corresponding functional vascular dysfunction, with evidence of reduced cerebral blood flow and brain volume. These structural and functional vascular changes corresponded with a decline in performance in the novel object recognition (NOR) test over time. None of these changes were seen in female DIO mice. Together our findings support the role of obesity-driven cerebrovascular dysfunction in the development of cognitive impairments. This work reinforces the critical importance of mixed-sex cohorts in metabolic research and identifies the cerebral microvasculature as a possible translational target for therapeutic intervention.

## Results

To quantify the effects of diet-induced obesity on cerebrovascular structure and function, and any associated cognitive changes, we combined longitudinal behavioural testing and MRI with endpoint vascular labelling, tissue clearing and three-dimensional light-sheet fluorescence microscopy in a 22-week mixed-sex DIO mouse model (Fig. 1A).

**Figure 1.**
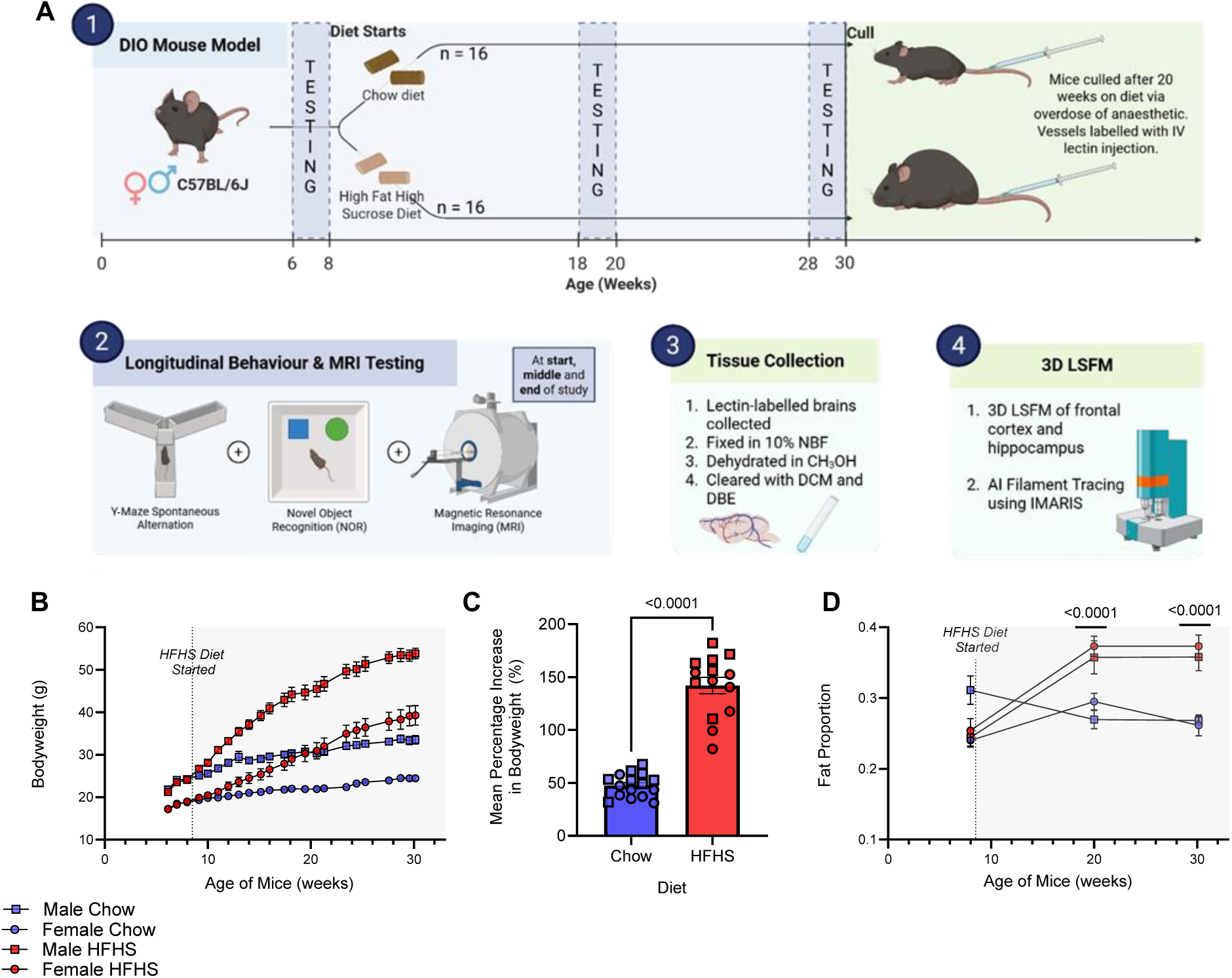
Study Design and Metabolic Metrics. (A1) Male and female C57BL/6 mice were randomized to receive high-fat high-sucrose (HFHS) diet (58% fat, 13% sucrose, DIO) or standard chow for 22 weeks. (A2) Longitudinal behavioural (Y-maze spontaneous alternation and novel object recognition (NOR)) testing and MRI scans were conducted at the beginning, middle and end of the testing period. (A3-4) Lectin-labelled brains underwent tissue clearing and subsequent 3D vascular imaging. (B) Mean bodyweight of Chow vs DIO mice over time, split by sex. Data collected from twice weekly weight measurements. Males and females analysed separately, via two-way repeated measures ANOVA (diet x time). (C) Mean increase in total bodyweight (%) in each diet group since the beginning of the study. Sexes were grouped, and analysed using unpaired t-test with Welch’s correction. (D) Mean bodyfat proportion of Chow vs DIO mice over the start, midpoint and end of the study, split by sex. Fat proportion calculated from MRI scans. Analysed via three-way repeated measures ANOVA with Tukey’s multiple comparisons test. The initial cohort comprised n = 8 mice per sex and diet. For B, C and D, the analysed cohort comprised *n* = 8 chow-fed and *n* = 7 HFHS-fed mice per sex. Data are presented as mean ± standard error of the mean. DIO, diet-induced obesity; IV, intravenous; MRI, magnetic resonance imaging; NBF, neutral buffered formalin; DCM, dichloromethane; DBE, dibenzyl ether; LSFM, light-sheet fluorescent microscopy; AI, artificial intelligence. Schematic created in BioRender.com.

### High-fat high-sucrose feeding induces obesity and increased adiposity in both sexes

To monitor development of the DIO phenotype, longitudinal measures of body weight and composition were taken. Mice fed a high-fat high-sucrose (HFHS) diet gained significantly more weight than chow-fed controls in both sexes (Fig. 1B), with DIO mice showing an approximately three-fold greater percentage weight gain over the course of the study (Fig. 1C). MRI-derived measures of body fat proportion confirmed increased adiposity in HFHS-fed mice at 10 weeks and again at the study endpoint after 22 weeks of diet exposure (Fig. 1D). Thus, the HFHS protocol produced a robust obese phenotype in mice of both sexes.

### Diet-induced obesity induces male-specific capillary rarefaction in the frontal cortex

To quantify structural cerebrovascular changes, at study endpoint, blood vessels were labelled with intravenous fluorescent lectin, and right hemispheres were optically cleared and imaged by 3D LSFM. Vascular networks were reconstructed within select regions using AI-assisted 3D filament tracing (Fig. 2A). Analyses were focused on the frontal cortex and hippocampus, two key regions for cognitive functioning which are consistently reported to be affected by both metabolic disease and AD^19–21^.

**Figure 2.**
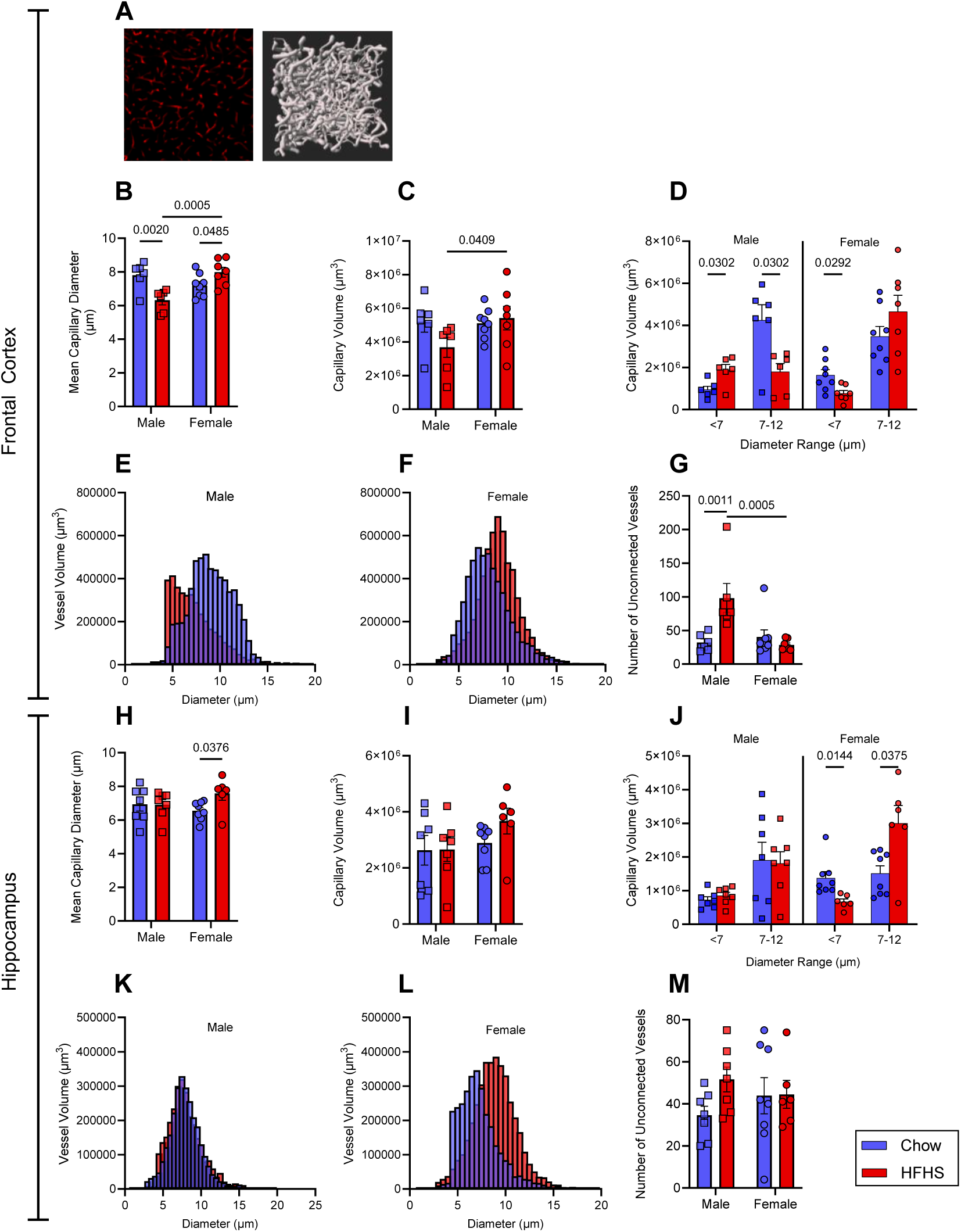
Structural Capillary Changes in the Frontal Cortex and Hippocampus of DIO Mice. (A) Representative lectin-labelled image and three-dimensional vascular reconstruction generated using IMARIS (v10.0) filament tracer. (B) Mean capillary diameter in the FC of DIO mice vs controls. (C) Total capillary volume in the FC of DIO mice vs controls. (D) Comparison of volume of small and medium frontal cortex blood vessels in DIO mice vs controls. (E-F) Histograms comparing the distribution of vessel volume across vessel diameters in male (E) and female (F) DIO and control mice. Diameter is split across bins of 0.5 μm width. (G) Number of unconnected vessels in the FC of DIO mice vs controls. (H) Mean capillary diameter in the HC of DIO mice vs controls. (I) Total capillary volume in the HC of DIO mice vs controls. (J) Comparison of volume of small and medium hippocampus blood vessels in DIO mice vs controls. (K-L) Histograms comparing the distribution of vessel volume across different vessel diameters in male (K) and female (L) DIO mice vs controls. Diameter is split across bins of 0.5 μm width. E) Number of unconnected vessels in the HC of DIO mice vs controls. Frontal cortex: n = 6 male chow, 6 male HFHS, 8 female chow and 7 female HFHS mice. Hippocampus: n = 7 male chow, 7 male HFHS, 8 female chow and 6 female HFHS mice. Capillaries were defined as vessels <12 μm in diameter; G and M quantify unconnected vessels in the unfiltered reconstructions. All bar charts presented as mean ± SEM. Data in B, C, G, H, I, and M analysed via two-way ANOVA with Fisher’s LSD. D and J analysed via unpaired t-tests, males and females analysed separately. ‘Small’ and ‘medium’ defined as diameters of <7 μm and 7-12 μm in diameter, respectively. FC, frontal cortex; HC, hippocampus, HFHS, high fat high sucrose; DIO, diet-induced obesity.

In the frontal cortex, male DIO mice exhibited a decrease in mean capillary diameter (capillaries defined as vessels with diameters below 12 µm) compared to chow-fed male controls, whereas female DIO mice exhibited an increase in mean capillary diameter relative to chow-fed females (Fig. 2B). The difference between male and female DIO, but not chow-fed, mice were also statistically significant, suggesting a sex-specific cerebrovascular response to metabolic stress. A significant sex × diet interaction supported these differing responses to diet.

Given previous evidence of a shift towards smaller cerebral vessels in obese mice^22^, we stratified capillaries into ‘small’ (< 7µm) and ‘medium’ (7-12 µm) diameter classes, to better characterise the diameter distribution. In male DIO mice, total capillary volume across both capillary classes was numerically lower than in chow-fed controls, although the difference did not reach statistical significance (p = 0.0816, Fig. 2C). The distribution of capillary volume was shifted towards smaller diameters (Fig. 2E) with an increased volume of ‘small’ capillaries, and a decreased volume of ‘medium’ capillaries (Fig. 2D).

Male DIO mice also had more unconnected vessel segments than chow-fed controls, consistent with greater fragmentation of the reconstructed vascular network (Fig. 2G)

In contrast, female DIO mice maintained overall capillary volumes compared to controls (Fig. 2C), but their distribution profiles shifted to the right, i.e. towards larger vessels (Fig. 2F), with a reduced volume of ‘small’ capillaries but preservation of ‘medium’ capillary volume (Fig. 2D). The difference in total capillary volume between male and female DIO mice was significant, whilst there was no difference between the total capillary volume of male and female chow mice (Fig. 2C).

Neither sex displayed differences in overall capillary length or number of branch points between DIO and controls (Supplementary. Fig. 1A-B).

These findings demonstrate a sex-dependent vascular response to the obese phenotype.

### Hippocampal vascular architecture is preserved in DIO males but displays capillary widening in females

We next asked whether similar microvascular changes occurred in the hippocampus. In male DIO mice no significant changes in hippocampal capillary diameter or volume were observed (Fig. 2H, I). In contrast, female DIO mice showed an increase in mean hippocampal capillary diameter (Fig. 2H), alongside a rightward shift in capillary volume distribution (Fig. 2J, L), consistent with the pattern observed in the frontal cortex. However, their total hippocampal capillary volume remained unchanged (Fig. 2I).

Neither sex displayed differences in overall capillary length or number of branch points between DIO and controls (Supplementary. Fig. 1C-D).

These data suggest regional vulnerability of the male cerebrovasculature to obesity, with a higher susceptibility in the frontal cortex than the hippocampus over the studied timescale. The capillary widening in females was observed across both brain regions, indicating a possible broader vascular compensation/adaptation to DIO in females.

### Bodyweight and adiposity are associated with frontal cortex capillary morphology in male mice

Next, to determine whether the severity of structural rarefaction was associated with the severity of the metabolic phenotype, we examined the relationship between bodyweight, bodyfat proportion and capillary structure in the frontal cortex.

In male mice, body weight negatively correlated with the mean capillary diameter, although the association with bodyfat proportion was non-significant (Fig. 3A-B). Neither body weight nor fat proportion was significantly associated with mean capillary diameter in females (Fig. 3C-D). Body weight and fat also correlated with the volume of small capillaries in males but not females (Fig. 3E-H). Thus, increasing body weight and adiposity in males was associated not only with narrower capillaries but also with a greater contribution of the smallest vessels to the microvessel network.

**Figure 3.**
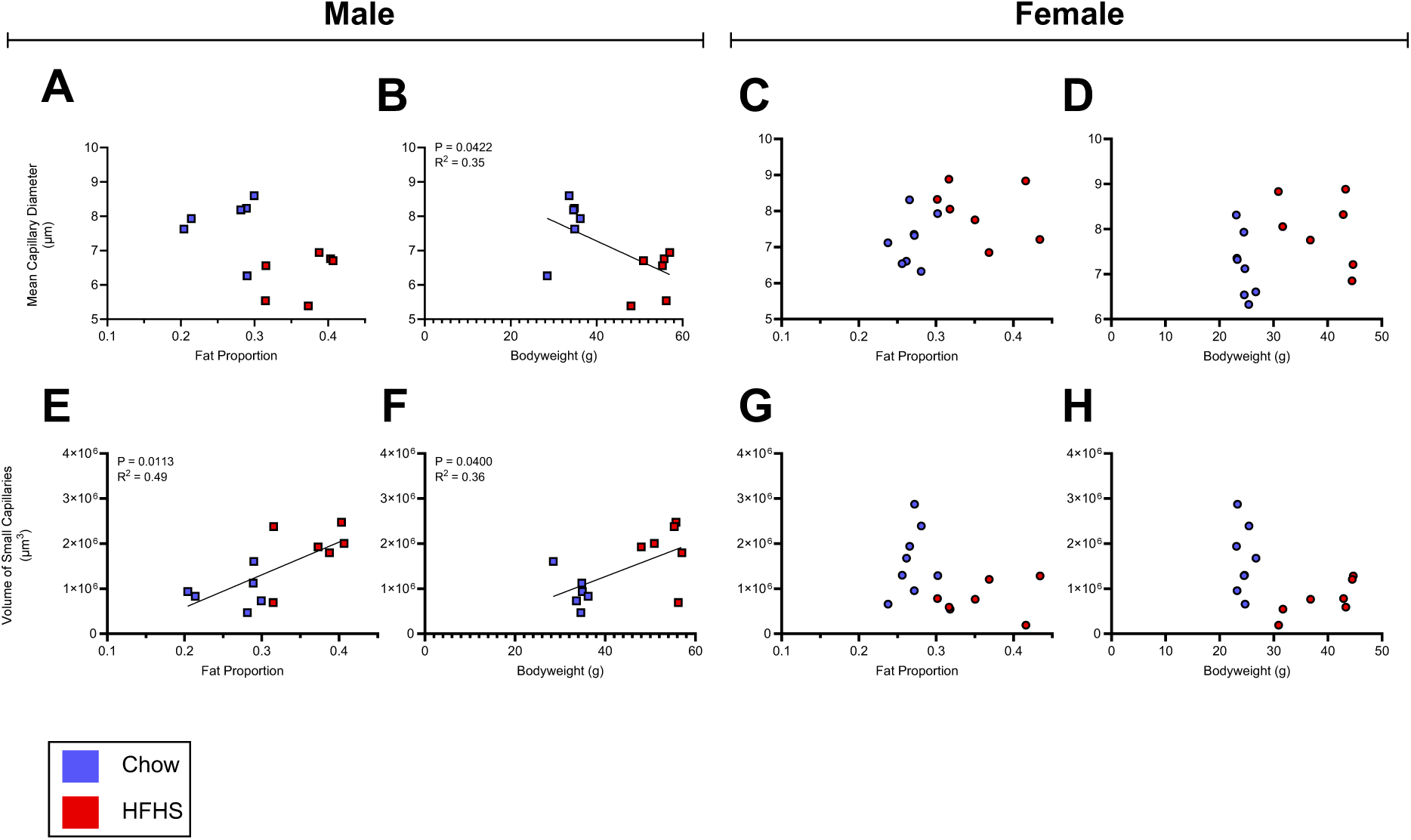
Sex-specific Correlations Between Weight, Fat and Structural Vascular Metrics in the Frontal Cortex. (A-B) Correlation between final bodyfat proportion (A) and final bodyweight (B) and mean capillary diameter in the frontal cortex of male mice, irrespective of diet group. (C-D) Correlation between final bodyfat proportion (C) and final bodyweight (D) and mean capillary diameter in the frontal cortex of female mice, irrespective of diet group. (E-F) Correlation between final bodyfat proportion (E) and final bodyweight (F) and small capillary volume in the frontal cortex of male mice, irrespective of diet group. (G-H) Correlation between final bodyfat proportion (G) and final bodyweight (H) and small capillary volume in the frontal cortex of female mice, irrespective of diet group. Sample size n=12 (males) and n=15 (females). Small capillaries are defined as vessels less than 7 μm diameter and capillaries are defined as less than 12 μm in diameter. Final bodyfat measurements were calculated from the endpoint MRI measurement after 22 weeks on diet protocols. Data presented as individual values with simple linear regression analyses plotted in B, E and F.

### Dynamic contrast-enhanced MRI reveals reduced frontal cortex blood volume in male DIO mice

To determine whether the structural capillary changes were accompanied by altered regional blood volume, we performed longitudinal, dynamic contrast-enhanced MRI (DCE-MRI) at baseline, midpoint and endpoint. Image stacks were registered to the Allen Mouse Brain Atlas, and region-specific gadolinium concentration was quantified (Fig. 4A). Regional gadolinium concentration was used as a proxy for blood volume. Measurements were normalised for region size and brain volume, to account for differences in tissue volume.

**Figure 4.**
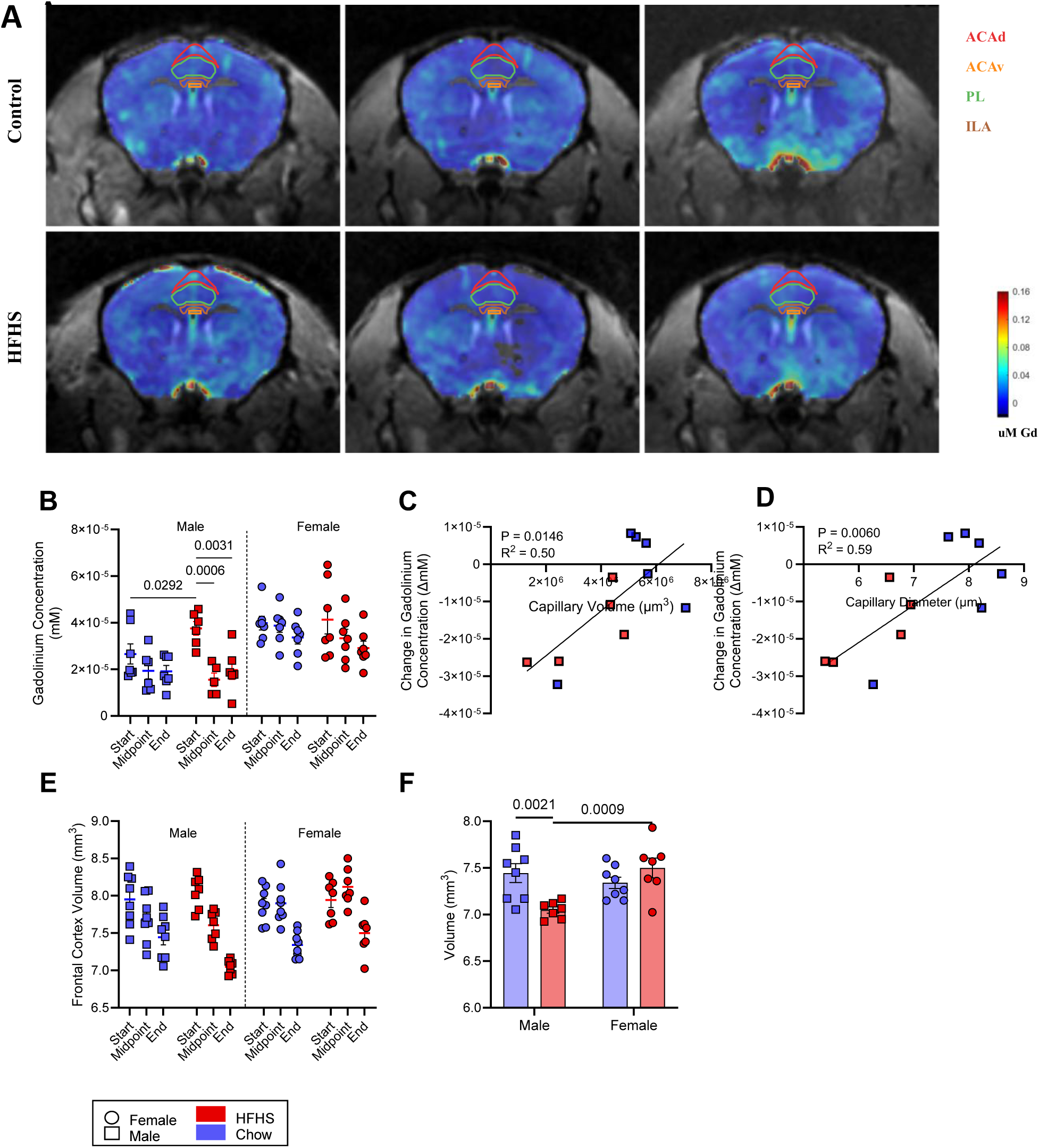
Blood Volume Changes in the Frontal Cortex of DIO Mice and Correlations with Capillary Structure. (A) Representative magnetic resonance imaging (MRI) maps of gadolinium (Gd) concentration in chow-fed (top) and high-fat, high-sucrose (HFHS)-fed (bottom) mice at baseline, midpoint and endpoint (left to right). Allen mouse atlas regions used to define the frontal cortex are outlined; the colour scale indicates Gd concentration in μM. (B) Frontal cortex Gd concentration across study timepoints, used as a proxy for blood volume. Analysed via mixed-effects model with Tukey’s multiple comparisons. (C, D) Correlations between change in Gd concentration and endpoint capillary volume (C) or mean diameter (D) in males, analysed vis simple linear regression. Change was calculated as endpoint minus baseline. (E) Frontal cortex tissue volume across study timepoints. Analysed via three-way ANOVA with Sidak’s multiple comparisons test (F) Endpoint frontal cortex tissue volumes, analysed via two-way ANOVA with Fisher’s LSD multiple comparison. Group summaries show mean ± standard error of the mean. Sample sizes: B: 6 HFHS males, 7 chow males,7 HFHS females, 7 chow females; C and D: 6 chow males, 5 HFHS males; E and F: 7 HFHS males, 7 HFHS females, 8 chow males, 8 chow females. ACAd and ACAv, dorsal and ventral parts of the anterior cingulate area, respectively; PL, prelimbic area; ILA, infralimbic area.

A difference in the mean baseline frontal cortex gadolinium concentration was seen between chow and DIO male mice (Fig.4A). Between baseline and 22 weeks on the HFHS diet, frontal cortex gadolinium concentration decreased in male DIO mice. No significant changes were detected over the same period in the male control mice, or either female group (Fig. 4B).

In the male mice, the change in frontal cortex gadolinium concentration was correlated with mean capillary volume and capillary diameter (Fig. 4C, D), linking the MRI-derived measures of vascular function with the 3D LSFM measures of capillary structure. No significant correlation was detected in the female mice, and no changes in blood volume were seen in the hippocampus of any group (data not shown).

Together, these findings indicate that diet-induced obesity is associated with a male-specific reduction in frontal cortex blood volume and a male-specific relationship between blood volume change and capillary structure.

### Structural MRI reveals frontal cortex volume loss in male DIO mice

We next examined whether the DIO phenotype was accompanied by changes to regional brain tissue volume. Structural MRI scans acquired at study baseline, midpoint and endpoint were registered to the Allen Mouse Brain Atlas and used to derive region-specific brain volumes. Frontal cortex volume decreased over time across all cohorts, consistent with an age-related reduction in volume. This decline was significantly greater in male DIO mice, supported by a significant sex x diet x timepoint interaction in a three-way ANOVA test (Fig. 4E). By the end of the study, the mean volume of the frontal cortex in male DIO mice was reduced compared to chow-fed males and DIO females (Fig. 4F).

Examination of other regions of the atlas showed that these volume changes were not reflective of a brain-wide volume change, but rather were region specific. There were no volume changes detected in the hippocampus (data not shown).

### Recognition memory declines selectively in male DIO mice and is associated with adiposity

To determine whether these structural and functional cerebrovascular changes were accompanied by cognitive impairment, mice underwent longitudinal, behavioural testing, including novel object recognition test and Y-maze spontaneous alternation.

No significant differences were observed in NOR performance between groups at baseline. After 10 weeks on the HFHS diet, male DIO mice displayed a reduction in NOR preference index (PI) relative to their own baseline performance, indicating impaired object recognition memory. This deficit persisted at 20 weeks, with male DIO mice continuing to perform worse than baseline (Fig. 5A). In contrast, female DIO mice did not exhibit a significant change in NOR performance relative to individual baselines and did not differ from female controls over time. At both midpoint and endpoint testing, body fat proportion correlated with NOR performance in male mice (Fig. 5B), such that greater adiposity was associated with poorer task performance. Of note, this relationship was not observed in females (Fig. 5C).

**Figure 5.**
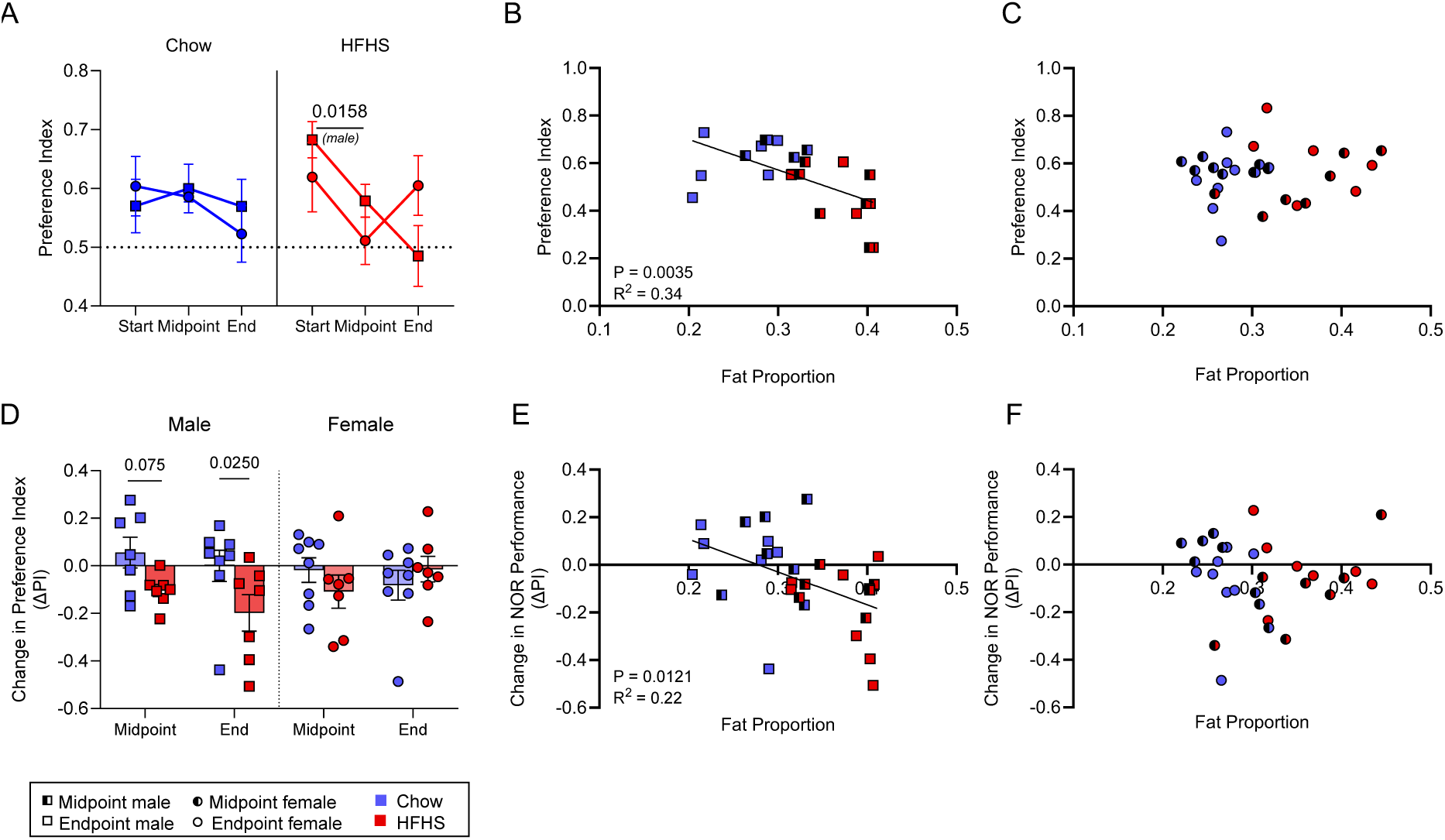
Cognitive Performance in NOR Test and Association with Bodyfat. (A) Novel object preference index of DIO mice vs controls across the study duration, split by sex. Data analysed via three-way ANOVA with Tukey’s multiple comparisons test. (B-C) Correlation between body fat proportion and NOR performance at the midpoint and endpoint of the study, irrespective of diet, in (B) males and (C) females. (D) Mean change in preference index over time. Measured as the difference between baseline NOR measurement and midpoint or endpoint NOR measure. Analysed via two-way ANOVA with Fisher’s LSD test. (E-F) Correlation between body fat proportion and change in NOR performance at the midpoint and endpoint of the study, irrespective of diet, in (E) males and (F) females. Sample size; 7 HFHS males, 7 chow males, 7 HFHS females, 8 chow females. ‘Start’ refers to the study baseline measurements taken at 6 weeks old, prior to diet protocols beginning. ‘Midpoint’ refers to the testing block occurring after 10 weeks on diet protocols. ‘End’ refers to the final study block, taken after 22 weeks on diet protocols. Dotted line in (A) indicates the PI value (0.5) where equal time is spent exploring the novel and familiar objects. Change in NOR performance is measured as the difference between baseline performance measured at study start and the timepoint of interest (either midpoint, after 10 weeks, or endpoint, after 22 weeks). All correlations presented as individual values and analysed via simple linear regression. PI, preference index; NOR, novel object recognition; HFHS, high fat high sucrose.

To quantify change in recognition memory over the course of the study, change in preference index (ΔPI) was calculated relative to each animal’s baseline. Male DIO mice displayed a decline in performance over time compared to chow-fed controls, indicating a decline in recognition memory over time (Fig. 5D). No such effect was seen in females.

To examine whether individual differences in adiposity were associated with recognition-memory performance, we correlated body fat fraction to PI and ΔPI. In males, greater adiposity was associated with poorer NOR performance and greater decline from baseline at both midpoint and endpoint assessments (Fig. 5B, E). Neither association was statistically significant in females (Fig. 5C, F).

### Spatial working memory is preserved in DIO mice

To assess whether obesity affected additional cognitive domains, the Y-maze spontaneous alternation task was used alongside the NOR test. No significant differences were seen in the spontaneous alternation behaviour of mice in the Y-maze at any time point (data not shown). These findings suggest that, within the timeline of this study, the cognitive impact of diet-induced obesity was selective (in this case for recognition memory).

### Frontal cortex vascular measures correlate with cognitive performance

Finally, we examined the relationship between structural and functional vascular measures in the frontal cortex and behavioural performance across the study. Endpoint PI correlated with structural capillary metrics including diameter and volume across both sexes, such that mice with narrower frontal cortex capillaries performed worse in the NOR test (Fig. 6A, B). ΔPI also correlated with capillary and diameter across both sexes, meaning that mice with narrower capillaries exhibited greater decline in object recognition memory over time (Fig. 6C, D).

**Figure 6.**
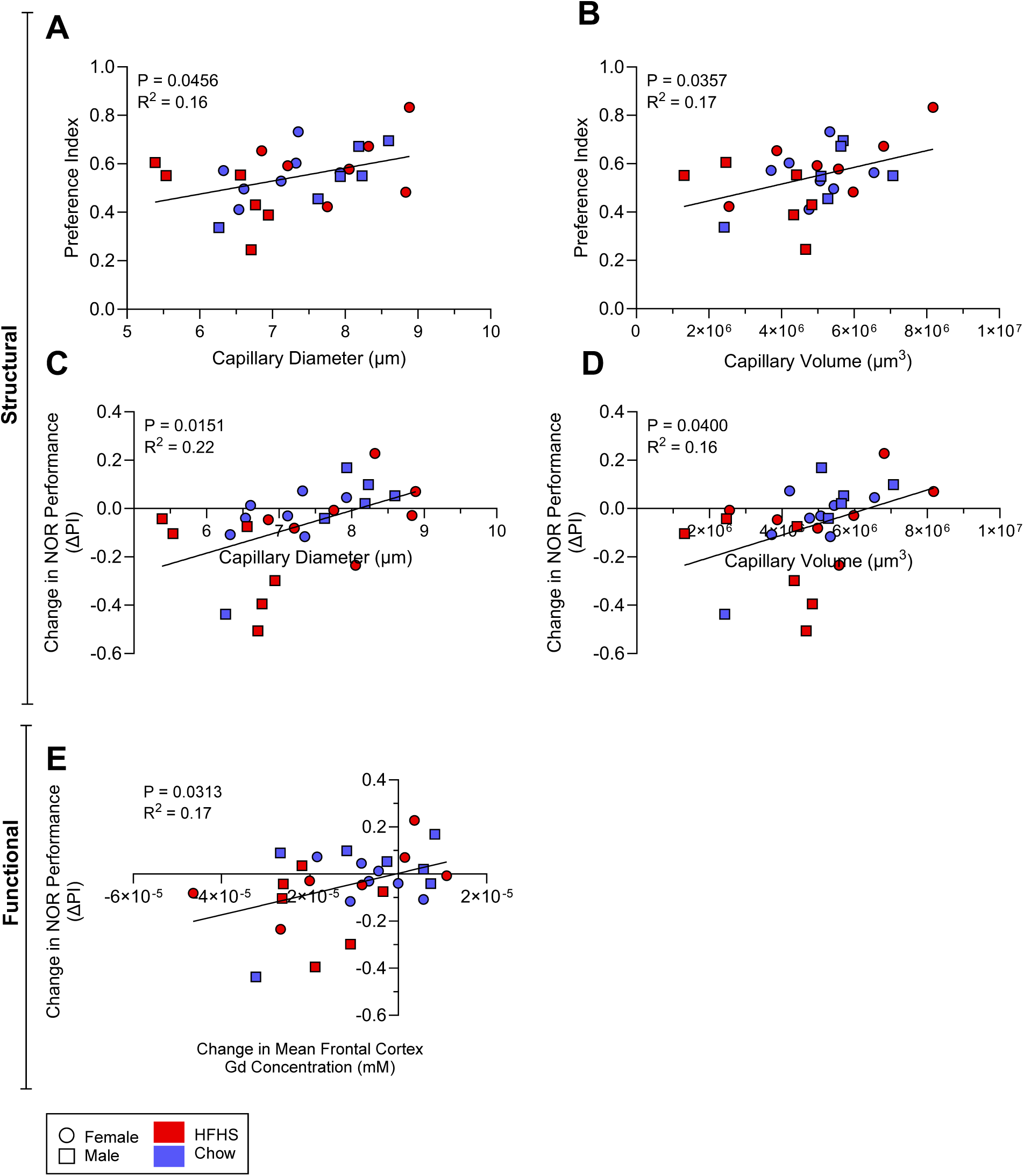
Correlation of Behavioural Performance with Vascular Structure and Vascular Function. (A-B) Correlation between raw NOR performance in the final behaviour testing week and mean capillary diameter (A) and total capillary volume (B) in the frontal cortex, irrespective of diet or sex. (C-D) Correlation between change in NOR performance over time and mean capillary diameter (C) and total capillary volume (D) in the frontal cortex, irrespective of diet or sex. (E) Correlation between change in NOR performance over time and change in gadolinium concentration in the frontal cortex, irrespective of diet or sex. Sample size; 7 HFHS females, 7 chow females, 7 HFHS males, 7 chow males. All data presented as individual values and analysed via simple linear regression. Change in NOR performance is measured as the difference in preference index from baseline measures taken at the start of the study and final measures taken at the end of the study after 22 weeks. Change in gadolinium concentration measured as the difference in gadolinium concentration taken from MRI scans at baseline and at the end of the study after 22 weeks. Capillary defined as a vessel with diameter <12μn. Gd, gadolinium; HFHS, high fat high sucrose; PI, preference index.

In males, ΔPI also correlated with the change in MRI-derived measures of frontal cortex blood volume (Fig. 6E). The larger the decrease in blood volume, the larger the decline in NOR performance.

Together these analyses link obesity-associated frontal cortex microvascular remodelling to memory impairments in male mice, highlighting a striking sex-specific vulnerability. A summary of the structural, functional and behavioural changes observed in the frontal cortex of DIO mice can be found in Table 1.

**Table 1.** Summary of Key Vascular Structural, Functional and Behavioural Changes Observed in the Frontal Cortex of DIO Mice. All measures taken from final endpoint measures, after 22 weeks on a HFHS diet. Changes are described relative to chow-fed, age-matched, same-sex controls. Changes reported as a decrease (↓), increase (↑), distribution shift to the left (←) or to the right (→) or no significant change (-). Symbols in brackets represent changes which did not reach the significance threshold of p<0.05 but were commented on in the main text.

|  | Metric | Additional Context | Change in DIO Male | Change in DIO Female |
|---|---|---|---|---|
| **3D LSFM:**<br>Structural changes | Mean Capillary Diameter |  | ↓ | ↑ |
|  | Total Capillary Volume |  | (↓) | - |
|  | Total Capillary Length |  | - | - |
|  | Number of Branch Points |  | - | - |
|  | Number of Unconnected Vessels | *Indicates how fragmented the vessel network is.* | ↑ | - |
|  | Vessel Diameter Distribution | *Indicates the most common vessel diameter. A shift to the left means vessels become thinner.* | ← | → |
| **MRI:**<br>Functional changes | Brain Volume |  | ↓ | - |
|  | Concentration of Gadolinium | *A proxy for the volume of blood in the region measured.* | ↓ | - |
| **BEHAVIOURAL TESTING:**<br>Cognitive changes | NOR performance (PI) | *A measure of object recognition memory. A higher value means better memory.* | ↓ | - |
|  | Change in NOR Performance (ΔPI) | *A measure of change in cognitive ability over time. Decrease indicates a worsening memory over time.* | ↓ | - |
|  | Y-maze Performance | *A measure of short term and spatial memory. A higher value means better memory.* | - | - |

## Discussion

Our integration of whole-brain 3D microscopy with longitudinal MRI and cognitive testing provides novel evidence that obesity induces a sex-specific cerebrovascular phenotype, supporting our hypothesis that obesity leads to microvascular changes in the development of early cognitive impairment. This approach extends previous descriptions of obesity-associated vascular pathology by examining structural, haemodynamic and cognitive measures together within individual animals, while also assessing sex differences. In male mice, the development of obesity was correlated with distinct structural capillary changes in the frontal cortex characterised by reduced capillary diameter, a marked distribution shift towards smaller vessels and a more fragmented vascular network. These structural changes were correlated with markers of functional changes in the microvasculature, including a decrease in frontal cortex blood volume and reduced frontal cortex tissue volume.

The decline in gadolinium concentration seen in the frontal cortex of obese male mice, used here as a proxy for reduced blood volume, complements previous clinical and preclinical studies showing reduced cerebral blood volume and perfusion in both obesity and Alzheimer’s disease^9,23^. In this study, its association with frontal-cortex capillary diameter and volume suggests that the development of structural cerebrovascular rarefaction may contribute to this. The reduced perfusion could explain the lower frontal cortex blood volume in obese males, which would be in keeping with human MRI studies showing that decreased CBF associates with brain atrophy^24^.

The impact on cerebral blood flow and blood-brain barrier (BBB) permeability, however, was not measured in this study. Given the growing recognition of BBB dysfunction as an early feature of cognitive impairment and neurodegenerative disease^25–27^, it would be valuable for future work to assess BBB permeability alongside vascular structure and blood volume, to understand whether barrier dysfunction accompanies these changes and is associated with recognition-memory impairment.

These markers of cerebrovascular dysfunction in DIO males were associated with worse performance in NOR tests, alongside a decrease in cognitive performance over time. While these associations do not establish causality, they do support a close relationship between frontal cortical vascular integrity and cognitive performance in this model. In our model, memory impairment was relatively mild. Clinical studies have demonstrated that mild cognitive impairment usually precedes the development of overt vascular dementia, suggesting the changes observed here may represent an early disease stage ^27^.

The regional specificity of the phenotype is also of note. In males, the frontal cortex and not hippocampus was affected. This contrasts with much of the existing literature, in which hippocampal changes are often reported alongside the frontal cortex in cognitive decline. It is possible that the frontal cortex is affected earlier in the disease progression, with hippocampal vascular dysfunction occurring later. Indeed, longer studies than this have reported hippocampal microvascular rarefaction occurring only after an extended exposure to metabolic stress and obesity ^9^. Another possibility is that the hippocampal response is more spatially heterogeneous and therefore not captured in the ROI approach used here. Regardless, our current data suggest that early obesity-associated vascular pathology is not uniformly distributed across the brain and that the frontal cortex may be a particularly vulnerable region in this model.

In contrast, female DIO mice did not display structural capillary rarefaction in either brain region. Instead, they exhibited increased capillary diameter and a distribution shift towards larger vessels across both frontal cortex and hippocampus, whilst maintaining capillary volume and without any evidence of impaired blood flow or cognitive impairment. These sex differences in the cerebrovascular profile and cognitive impact of obesity were an especially striking finding. Many *in vivo* pre-clinical studies still rely predominantly on studying male animals so that these sex differences may be overlooked. In humans, sex differences exist but with women being reported to have a higher susceptibility to many forms of dementia, including Alzheimer’s disease ^28^. This may be explained, at least in part, by increased longevity in females and thus greater risk of developing age-related neurodegenerative diseases^29^. In post-menopausal women there may also be effects from hormonal transition. In keeping with this, reproductive hormones in pre-menopausal women may convey a degree of neurovascular protection ^30^. Oestrogen, in particular, moderates vasodilation via nitric oxide ^31^, and its loss following menopause is thought to be a potential risk factor for neurodegenerative disease ^28,29^. Consistent with this, studies have shown that young, ovariectomised, female mice and rats display increased BBB permeability compared to controls with intact reproductive systems ^32,33^, and greater cognitive decline is reported in women with early-onset menopause ^28^. There is also evidence that females are protected against microvascular rarefaction in other disease pathologies^34–37^.

The results of the present study fit well within this narrative, as the focus here is on the early development of vascular-associated cognitive decline in obesity, rather than overt dementia and late-stage neurodegeneration. It is therefore likely that sex modifies cerebrovascular risk in metabolic disease in an age-dependent manner. Earlier in life, females may retain a degree of vascular protection, possibly mediated by reproductive hormones, which helps preserve endothelial function and BBB integrity in obesity ^38^. With ageing, this protection may diminish, increasing female vulnerability in older human cohorts.

In this context, the vascular preservation observed in the female DIO mice here may reflect an early, hormone-supported compensatory response to metabolic stress. The larger capillary diameters in both the frontal cortex and hippocampus of these females are consistent with this interpretation, and suggests that females may compensate against obesity-driven microvascular rarefaction through adaptive vasodilation. Whether this is just a brain-wide response, or reflective of a more systemic multi-organ phenotype, is yet to be determined, but it provides a possible explanation for why female DIO mice were relatively protected from the vascular and behavioural changes seen in DIO males.

Interestingly, the overarching relationship between diet-induced obesity, vascular dysfunction and cognitive decline appears to be comprised of both sex-dependent and sex-independent mechanisms. The promotion of frontal cortex microvascular rarefaction and reduced cerebral blood volume by a HFHS diet (associated with body fat proportion) was male specific. Yet, the association between vascular parameters and cognitive performance was not obviously restricted to one sex. In practice, this meant that females appeared to be protected against obesity-associated cerebrovascular rarefaction, but that variation in vascular integrity, when present, impacted cognitive performance in a similar way across the cohort. This suggests that sex modifies susceptibility to obesity-induced vascular dysfunction more strongly than it modifies the downstream relationship between vascular dysfunction and cognition.

These associations do not establish that vascular changes cause cognitive decline, and endpoint measurements of capillary structure cannot determine whether such remodelling precedes memory impairment. Future work should focus on understanding the mechanisms underlying the observed changes.

However, taken together, these results provide evidence that microvascular dysfunction is an early and functionally relevant pathological feature linking metabolic disease to cognitive decline. They establish the critical role of vascular biology in obesity-associated cognitive impairment and underscore the importance of incorporating sex as a biological variable in studies of brain health in metabolic disease. Interventions aimed at preserving microvascular integrity may be effective in reducing obesity-associated cognitive decline, particularly if applied early and with attention to sex-specific biology.

## Methods

### Animals and Regulatory Approvals

All animal procedures were approved by the University of Cambridge Animal Welfare and Ethical Review Body (AWERB) and conducted according to the UK Animals (Scientific Procedures) Act 1986 (Amendment Regulations 2012), under Home Office project licence P160D4B75.

Male and female C57BL/6J mice were obtained from Charles River Laboratories, UK, at 5-6 weeks of age. Animals were housed in same-sex groups of four, in individually ventilated cages with standard bedding and enrichment, on a reverse light-dark cycle in temperature and humidity-controlled conditions. Access to food and water was provided on an *ad libitum* basis. Mice were given at least 7 days to habituate before study start. All animals were handled regularly, at a minimum of twice a week for the duration of studies, to reduce handling-related stress.

A total of 32 mice were included at study start, comprising 16 males and 16 females. Within each sex, 8 mice were assigned to the HFHS diet and 8 to standard chow. Two animals in the DIO group, one male and one female, were culled on welfare grounds during the study, resulting in a final cohort of 30 mice. Exact group sizes for each experiment are provided in figure labels.

### Diet-Induced Obesity Model

To induce obesity, mice were fed a high-fat high-sucrose (HFHS) diet (D12331, Research Diets Inc.). The diet provided 58% energy from fat, 25.5% from carbohydrates and 16.4% from protein, with 12.6% kcal from sucrose, for a total of 5558.5 kcal kg^-1^ of diet. Control mice received standard (SAFE® A03, Scientific Diets Inc.) which provided 3395 kcal kg^-1^, with 13.5% from fat and no added sucrose. Diets were provided *ad libitum*, and the HFHS diet was changed weekly to avoid spoilage.

### Experimental Study Design

The study used a longitudinal design with repeated behavioural and MRI assessments. Baseline measurements were obtained before diet allocation. Follow-up testing was then performed after 10 weeks on diet and again at the study endpoint after 20 weeks on diet. Each testing block consisted of behavioural testing in week 1, followed by an MRI scan in week 2. At the final time point, directly following the final MRI scan, animals underwent intravenous fluorescent lectin labelling of the vasculature and were then euthanized for tissue collection.

Behavioural testing was performed during the dark phase. The Y-maze was conducted on day 1 of the behavioural week, followed by a rest day, and then a 3-day novel object recognition task from day 3 to day 5. Animals were given 60 minutes to acclimatise to the testing room before each session. Males were tested before females, with a 60-minute interval between sexes to allow room ventilation and scent clearance. Within each sex, testing order was otherwise randomized.

### Metabolic Monitoring

Bodyweight was recorded twice weekly throughout the study. Body fat proportion was measured by MRI as described below.

### Novel Object Recognition

#### Apparatus

Novel object recognition was performed in a square arena with an infra-red transparent base and anti-reflective walls optimised for infra-red video tracking (Ugo Basile, Product Code 47432-IR). The arena was illuminated from below using an infra-red floor (LUX 100, Ugo Basile). Behaviour trials were recorded using an IR CMOS camera (Teledyne Blackfly GigE, BFLY-PGE-13H2M-CS) mounted above the arena. Live tracking was performed using EthovisionXT v17.5 (Noldus Information Technology).

#### Testing Protocol

The NOR test followed a three-day protocol. On day 1, mice were given 5 minutes to habituate to the empty arena. Twenty-four hours later, during the ‘familiarisation’ phase, two identical objects were placed equidistant in the arena. The mouse was placed in the centre of the arena, facing the north wall, and given 5 minutes to explore. On day 3, during the ‘testing’ phase, one familiar object from day 2 (a triplicate to avoid any scent-aided recognition), and a novel object were placed in the arena and the mouse again given five minutes to explore. Objects were fixed in place to the arena floor with adhesive Velcro, to prevent movement during exploration.

Objects were selected to be visually distinct but had similar heights and were constructed of materials that were not reflective to reduce any inherent preferences for a particular object. Testing during the dark phase helped with this ^39,40^.

For every trial, the object pairs were randomly permutated between position (L or R) in the arena, as well as which one was “novel” for each subject. All objects and the arena were cleaned with 70% ethanol spray between subject trials.

#### Behavioural Analysis

Trial videos were analysed in EthovisionXT (v17.5, Noldus Information Technology). Dynamic subtraction was used for mouse detection, with three-point tracking of nose point, body centre and tail base. Tracking began once all three body points were identified within the arena for a duration of five seconds, and continued for 5 minutes, after which the tracking automatically stopped.

Object exploration was defined as the total duration for which the nose point or body centre was touching the object, or when the nose point was inside of the object zone (a 2cm border around the object). Head direction towards an object was the total duration that the nose point was directed towards to object as determined by the direction of all three tracking points. Total distance travelled, mean velocity and time spent stationary were all also calculated.

The primary outcome measure was the preference index (PI), calculated as the proportion of total object exploration time spent exploring the novel object ^41^. Change in preference index over time (ΔPI) was calculated relative to each animal’s baseline performance.

### Y-maze spontaneous alternation

#### Apparatus

Spontaneous alternation testing was performed in the dark phase, in a Y-maze, with three arms equally spaced at 120° angles to the centre. Each arm was 35 cm long and 5 cm wide. The arena had an infra-red transparent base and walls made of dark, anti-reflective material optimised for infra-red video tracking (Ugo Basile, Product Code 40173-IR). The maze was illuminated from below using an infra-red floor (Ugo Basile, LUX 100). Trials were recorded and tracked using the same infrared camera and tracking software used in NOR testing.

#### Testing Protocol

Each mouse was placed into the centre of the maze and allowed to explore freely for 8 minutes. The maze was cleaned with 70% ethanol between trials.

#### Behavioural Analysis

Tracking of trials was performed in Ethovision XT. Dynamic subtraction was used to identify the mouse against the background, with three-point tracking of nose point, body centre and tail base. Tracking began once all three body points were detected within the arena for a duration of 5 seconds and lasted for a duration of 8 minutes.

An arm entry was defined as when all three body points had crossed the threshold of an arm. Alternations were defined as entries into all three arms consecutively without repeated arm visits. The primary outcome was the alternation index, calculated as the number of alternations divided by the total possible max alternations ^42^. Total arm entries, number of direct/indirect arm revisits, total distance travelled, mean velocity, and time spent stationary were also recorded.

### Magnetic Resonance Imaging

MRI was performed on a 9.4T horizontal bore preclinical MRI machine (Bruker BioSpec 94/20; Bruker Ltd.) running ParaVision software (v7.0). Prior to scanning, mice were anaesthetised via inhalation of isoflurane (1000mg/g, Iso-Vet®) and tail-vein cannulated with a 30G needle. Cannulas were flushed with heparinized saline (10 I.U/ml).

During scanning, mice were placed prone in the scanner, with the head stabilised with ear bars and the manufacturer standard mouse brain coil (4 channel array). Throughout imaging, respiratory rate and internal temperature were monitored continuously with a chest pad and rectal probe (SA Instruments; Stony Brook, NY), with a heated water system used to maintain body temperature. Anaesthesia was maintained with a constant flow of isoflurane and O_2_ via a nose cone (1-2% in 1 L/min O_2_) and dynamically adjusted to maintain the respiratory rate within the physiological range.

### Body Composition MRI

Body fat proportion was quantified using a 3-point Dixon method using the IDEAL multi-Dixon algorithm ^43^ (FLASH, TE=2.143, 2.3879, 2.6328ms; FOV 75×40×40mm, matrix 128×96×96 resolution 0.586×0.417×0.417mm) sequence and was calculated as the proportion of signal intensity from the mouse trunk that came from fat compared to water.

### Structural Brain MRI

Structural scans were performed for multiparametric mapping (MPM, with magnetisation transfer MTw, proton density PDw and T1w contrast). Post-injection, PD and T1w images were repeated to measure T1 changes. For MPM images had a 32×17.5×12.5mm^3^ with a matrix of 256×140×100 for an isotropic resolution of 125µm. Acceleration of 2.24× was achieved in each acquisition by zero-filling (factor 1.5 in each direction) and 1.5 partial Fourier in the slice direction. The flip angle was 6° for MTw and PDw, 30° for the T1w image. TR/TE1 was consistent at 25ms and 2.41ms for each of those.

### Dynamic contrast-enhanced MRI

Vessel perfusion and blood flow dynamics were characterised with a dynamic contrast enhancement scan with a fast low angle shot pulse sequence (DCE FLASH; TR/TE 35/1.5ms, flip angle 10°, with 100 frames acquired at 3.5 second intervals). The field of view was 20.00 x 20.00 x 8.1 mm^3^, with a voxel matrix of 96x96x9. The gadolinium bolus (Gadovist Bayer PLC, 50 µl of 0.1 mmol/ml) was administered via the cannula 30 seconds into the scan, delivered uniformly over a period of 8 seconds.

After scanning animals were placed into a clean, empty recovery cage in a heating cabinet with access to ad libitum water and food and once awake were returned to their home cage.

### MRI Analyses

Multi parametric mapping was performed following standard workflows described in ^44^.

Anatomical labelling used the Turone Mouse Brain Template and Atlas (TMBTA)^45^, with structure labels re-mapped onto the hierarchical ontology of the Allen Mouse Brain Common Coordinate Framework (CCFv3)^46^. Structures were examined at coarse and mid-levels of this hierarchy (designed for this study).

A study-specific template was constructed from the cohort using the ANTs template-construction pipeline^47^, and each animal’s images were registered to this template by symmetric normalisation (SyN), combining a nonlinear warp with an affine transform. The template was in turn aligned to the TMBTA atlas space, so that per-subject transforms (affine plus SyN warp field) mapped directly between native and atlas coordinates. Regional volumes were calculated by integrating the Jacobian determinants from the combined image registration transformations over each ROI.

Regional gadolinium concentration was estimated from the contrast-induced change in longitudinal relaxation rate (ΔR1), converted to concentration using an estimated relaxivity of Gadovist of 5 mM⁻¹s⁻¹ at 9.4 T.

### In-Vivo Vascular Labelling

At the final experimental time point, the vasculature was labelled using fluorescent lectin (100 µl, DyLight-649, Invitrogen, L32472), administered via the indwelling tail vein cannula immediately following the MRI scan whilst still under anaesthesia. After 5 minutes, to allow the lectin to circulate, euthanasia by overdose of sodium pentobarbital (Dolethal 200mg/ml, Vetoquinol UK) was performed, administered via the same tail cannula.

### Tissue Collection and Processing

Following confirmation of death, brains were rapidly dissected out and split sagitally into hemispheres. The right hemisphere was used for light-sheet fluorescence microscopy.

### Fixation and Dehydration for 3D LSFM

Right hemispheres were fixed in 10% neutral buffered formalin (NBF, Sigma-Aldrich for 36 hours at room temperature in the dark. Tissues were then dehydrated in increasing concentrations of methanol (20%, 40%, 60%, 80%, 100%) diluted with PBS as appropriate for a minimum of 5 hours per step, before being stored in a final concentration of 100% methanol (Sigma-Aldrich) in the dark at room temperature until clearing.

### Optical Clearing for 3D Imaging

Dehydrated right brain hemispheres were incubated in a solution of 66% dichloromethane (DCM) and 34% Methanol (Sigma Aldritch, US) for 180 minutes on an orbital shaker, before two further 30-minute incubations in 100% DCM (Sigma Aldrich, US). Tissues were then transferred into di-benzyl ether (DBE, Sigma Aldritch) and incubated overnight, before a final transfer into ethyl cinnamate (Eci, Sigma Aldritch, US), the scanning medium for the microscope.

### 3D Light-sheet fluorescent microscopy

Scanning was performed on the Miltenyi UltraMicroscope Blaze™ (Miltenyi Biotec, Germany) in conjunction with the ImSpector microscope controller software (v7.5.2, Abberior Instruments, Germany).

### Image Acquisition

Cleared brain hemispheres were imaged in ethyl cinnamate, using a 4x objective with 1.66x zoom, and a Z-step size of 3 µm. Lectin-labelled vessels were imaged with 630/30 excitation and 680/30 emission at 70% laser power and 50 milliseconds exposure. Tissue autofluorescence was also imaged with 560/40 excitation and 620/60 emission at 100% laser power and 50ms exposure to aid in anatomical navigation.

Two 16-bit tiff stacks were acquired per brain, one centred on the frontal cortex and one for the hippocampus. The field of view for all acquisitions was 2048 x 2048 voxels (the z length was variable between scans to ensure the desired region was captured), translating to 1996 x 1996 x 2000-3000 µm for the frontal cortex and 1996 x 1996 x 1000-2000 µm for the hippocampus. Navigation through the brain in the horizontal plane was assisted by the Paxinos and Franklin’s Mouse Brain Atlas (v5 2019) ^48^.

### Region Selection and AI Filament Tracing

Representative regions of interest (ROI) were selected in the frontal cortex and hippocampus light-sheet scans for 3D vascular reconstruction. ROI sizes were uniform across samples. In the frontal cortex, a ROI of 531 x 300 x 162 µm was placed to encompass the cingulate cortex and prelimbic cortex, at approximately Bregma -1.98mm. In the hippocampus, three ROIs were placed, each measuring 300 x 300 x 100 µm, in CA1, CA3 and the dentate gyrus at approximately Bregma -1.688mm.

Three-dimensional vessel maps were generated in IMARIS (v10.0, Oxford Instruments) using the AI filament tracing module. A minimum seed point diameter of 3 µm was used, with a threshold intensity of 1000, together with an iterative train and predict machine learning algorithm to identify vessel segments. Vessel statistics were automatically calculated in IMARIS.

### Capillary Filtering and Vascular Metrics

To focus analyses on the cerebral microvasculature, capillary-specific metrics were derived by filtering vessels according to diameter. The threshold diameter for a capillary classification was set at 12 µm. Vascular metrics extracted included total vessel volume, mean vessel diameter, total vessel length, number of branch points and number of unconnected vessel segments. Because ROI dimensions were consistent within each brain region, raw values rather than density-normalized values were used for comparison.

### Statistical Analyses

All statistical analyses were performed in GraphPad Prism 10.5.0 (GraphPad Software, USA). Data are presented as a mean ± SEM unless stated otherwise. Significance is defined as p < 0.05. Details of specific statistical tests used are laid out in figure legends or when results are described in the main text, where appropriate. Exact p values are included in their respective graphs. Sex differences in regression slopes were assessed using ANCOVA models including sex, the predictor and their interaction.

## Acknowledgements

We are grateful to the University Biomedical Services (UBS) at the Innes Building, University of Cambridge, for performing animal husbandry. We would also like to acknowledge staff at the Disease Model Core (DMC) Facility for assistance with animal and diet procurement. Work at the DMC is supported by the Medical Research Council [grant number MC_UU_00039] and the Wellcome Trust [grant number 226800/Z/22/Z]. We additionally thank novo employees Dr Camilla Ingvorsen for 3D LSFM support and Jeanette Bannebjerg for technical support with brain tissue clearing.

---

**Supplementary Figure 1.**
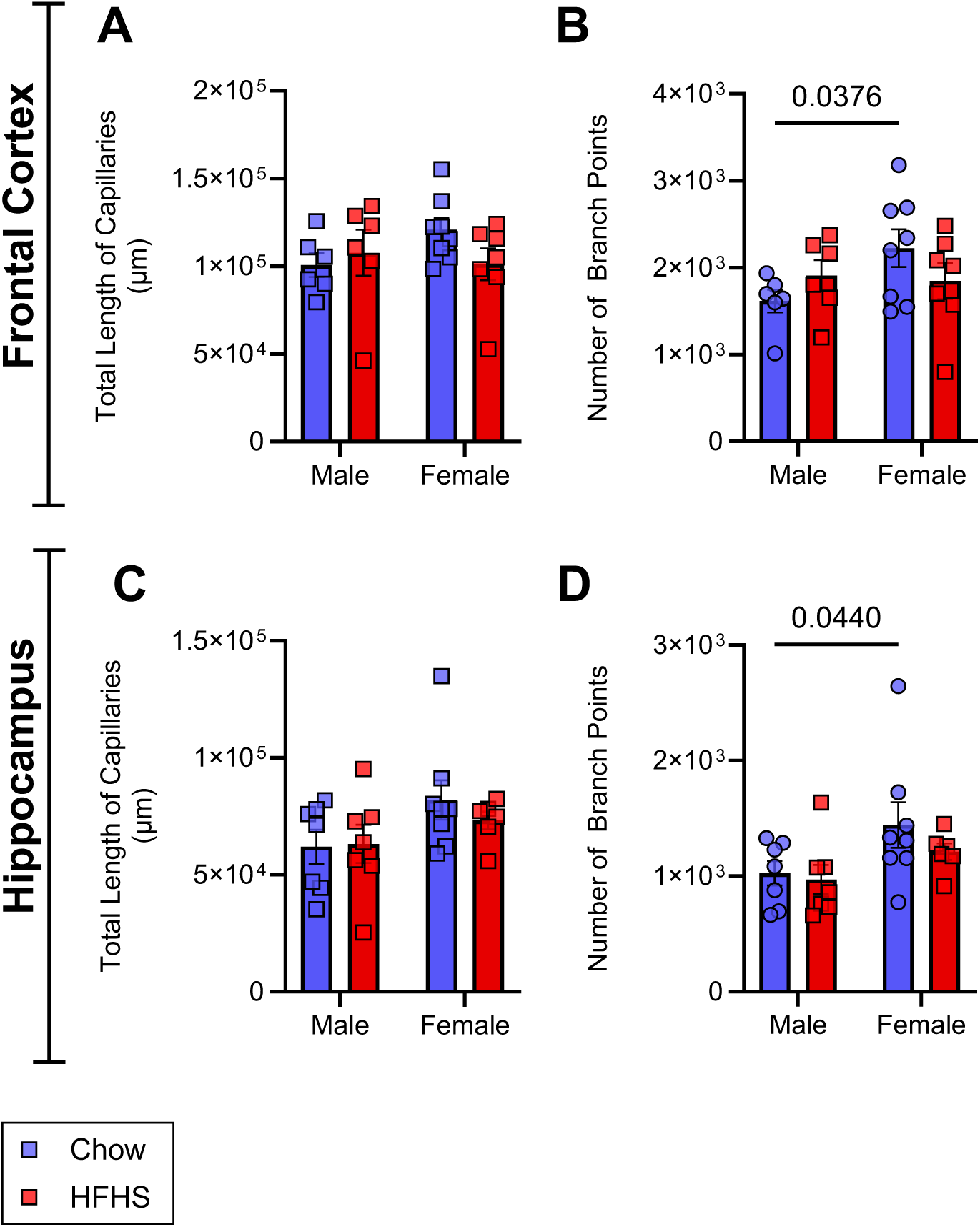
Additional Structural Capillary Metrics in the Frontal Cortex and Hippocampus of DIO Mice. (A) Total capillary length in the FC of DIO vs control mice. (B) Number of branch points in the FC of DIO vs control mice. (C) Total capillary length in the FC of DIO vs control mice. (D) Number of branch points in the FC of DIO vs control mice. Frontal cortex: n = 6 male chow, 6 male HFHS, 8 female chow and 7 female HFHS mice. Hippocampus: n = 7 male chow, 7 male HFHS, 8 female chow and 6 female HFHS mice. Capillaries were defined as vessels <12 μm in diameter. All bar charts presented as mean ± SEM. Data analysed via two-way ANOVA with Fisher’s LSD. FC, frontal cortex; HC, hippocampus, HFHS, high fat high sucrose; DIO, diet-induced obesity.

